# Aptamer-antibody chimera sensors for sensitive, rapid and reversible molecular detection in complex samples

**DOI:** 10.1101/2023.08.08.552518

**Authors:** Dehui Kong, Nicolo Maganzini, Ian A.P. Thompson, Michael Eisenstein, H. Tom Soh

## Abstract

The development of receptors suitable for the continuous detection of analytes in complex, interferent-rich samples remains challenging. Antibodies are highly sensitive but difficult to engineer in order to introduce signaling functionality, while aptamer switches are easy to construct but often yield only modest target sensitivity. We present here the programmable antibody and DNA aptamer switch (PANDAS), which combines the best features of both systems by using a nucleic acid tether to link an analyte-specific antibody to an internal strand-displacement (ISD)-based aptamer switch that recognizes the same target. The monoclonal antibody mediates initial analyte binding due to its higher affinity; the resulting increase in local analyte concentration then leads to cooperative binding and signaling by the ISD switch. We developed a PANDAS sensor for the clotting protein thrombin and show that this design achieves 100-fold enhanced sensitivity compared to using an aptamer alone. This design also exhibits reversible binding, enabling repeated measurements with temporal resolution of ∼10 minutes, and retains excellent sensitivity even in interferent-rich samples. With future development, this PANDAS approach could enable the adaptation of existing protein-binding aptamers with modest affinity into sensors that deliver excellent sensitivity and minute-scale resolution in minimally prepared biological specimens.

## Introduction

Most approaches to molecular detection are designed for single-timepoint measurements, but there is considerable value in being able to track changes in analyte concentration dynamically. This is particularly important when tracking clinically-relevant markers in biofluids, where continuous measurements can provide a real-time indicator of disease states^1,2^, metabolic activity^3^, hormonal dysfunction^4,5^, and other physiological processes^6,7^. However, it remains a challenge to develop simple, generalizable biosensor architectures that are suitable for continuous molecular detection. Monoclonal antibodies generally offer the best sensitivity and specificity, but conventional immunoassays such as enzyme-linked immunosorbent assays (ELISA) require the use of a secondary reporter antibody, and thus are limited to single-timepoint measurements^8,9^. One recently-described alternative approach converts antibodies into a molecular pendulum construct that achieves reversible binding-induced electrochemical signaling^10^. However, this approach relies on the antibody’s intrinsic properties and thus has limited capacity for tunability to optimize its sensitivity. In contrast to antibodies, it is relatively straightforward to engineer and tune nucleic acid-based aptamers to undergo a reversible conformation change upon target binding. When coupled with a reporter moiety such as a redox tag or fluorophore, aptamer-based molecular switches can be used to continuously measure target molecule concentrations directly in complex samples^11–15^. For example, the commonly used intramolecular strand displacement (ISD) design, in which signaling is inhibited in the absence of target by a complementary DNA strand that hybridizes to the aptamer’s target-binding domain, offers a generalizable approach to aptamer switch development^16,17^. Nevertheless, aptamer switch design approaches introduce important trade-offs, as competition-based switching mechanisms can greatly reduce aptamer affinity^18^. Given that aptamers generally exhibit lower target affinity than antibodies, this can render them insufficiently sensitive for real-world detection applications. While there are alternative strategies for designing structure-switching aptamers such as screening or truncation, these strategies all tend to yield aptamers with limited sensitivity that is generally inadequate for clinically relevant protein biomarkers, which often exhibit physiological concentrations in the sub-nanomolar range^19^.

In this work, we present a molecular switch design that leverages both the high affinity of antibodies and the molecular switch functionality that is accessible with aptamers. Our Programmable Antibody and DNA Aptamer Switch (PANDAS) design links a structure-switching aptamer to the Fc-region of a modified antibody through a short DNA scaffold. Signaling is achieved through an ISD aptamer^16,17^ design with a fluorescent readout. Normally, such sensors exhibit modest target affinity, but in the context of PANDAS, the affinity of the ISD aptamer is greatly enhanced through the effects of antibody-mediated cooperative binding. When the antibody binds the target molecule, the coupled ISD aptamer experiences a high local target concentration that promotes rapid binding and conformation change to generate a fluorescent signal. As a demonstration, we have generated a PANDAS-based sensor for the blood clotting protein thrombin. We show that this construct can achieve ∼100-fold enhanced thrombin affinity relative to the aptamer switch on its own. Importantly, the ISD design also confers the ability to tune sensor affinity by altering the length of the displacement strand, and we show that an optimally tuned version of this PANDAS can achieve an effective equilibrium dissociation constant (*K*_*D*_) of ∼100 pM with minimal background. Furthermore, this PANDAS construct binds reversibly, enabling us to obtain continuous measurements of changing thrombin concentrations with ∼10-minute temporal resolution. Finally, our sensor design maintains excellent target specificity, and exhibited robust sensitivity and temporal resolution even when tested in an interferent protein-rich sample matrix. Given the modular design of the PANDAS construct, which requires no meaningful engineering of either the antibody or aptamer sequence beyond the introduction of a linker domain and displacement strand sequence, we believe this strategy should be broadly applicable for repurposing low-affinity aptamer switches to detect low-abundance targets in complex samples.

## Results

### Rationale and design for an antibody-enhanced aptamer switch

The PANDAS reagent design takes advantage of the benefits of both aptamer switches and monoclonal antibodies. The aptamer component of the PANDAS construct makes use of the ISD design to achieve fluorescent signaling upon aptamer-target binding (**Figure 1a**). Here, the aptamer is linked to a short displacement strand (DS) domain that is complementary to a portion of the aptamer’s binding motif. The DS hybridizes with the aptamer in the absence of target but is displaced by ligand binding as the free target concentration increases. This leads to target-dependent conformational switching of the ISD aptamer that can be fluorescently measured using a fluorophore-quencher pair coupled to the ends of the construct. However, because the ISD design introduces an intramolecular competitor within the aptamer switch that interferes with binding, the resulting ISD construct will always exhibit reduced affinity and temporal resolution compared to the native aptamer. The PANDAS design compensates for this reduced affinity by coupling the ISD aptamer switch to an antibody that recognizes the same target through a different epitope with higher affinity (see **Supplementary Fig. S1** for a schematic of the assembly process). The result is a hybrid molecular sensor that achieves high sensitivity through cooperative binding (**Fig. 1b**). The high-affinity antibody binds the target at low concentrations at which the aptamer switch would not otherwise bind, and the resulting proximity between the aptamer and antibody-bound target induces aptamer binding, switching, and signaling at substantially lower concentrations than can be detected with the ISD alone. The ISD design also allows fine-tuning of the aptamer switch function through changes in its DNA sequence. For example, increasing the length of the DS will shift the equilibrium toward the fully folded and quenched state by forming a more stable stem structure^17^. This results in background signal suppression at the expense of decreased aptamer switch affinity and temporal resolution. Conversely, decreasing the DS length reduces the strength of intramolecular binding competition, thus increasing aptamer switch affinity and temporal resolution at the cost of a higher background signal.

**Figure 1.**
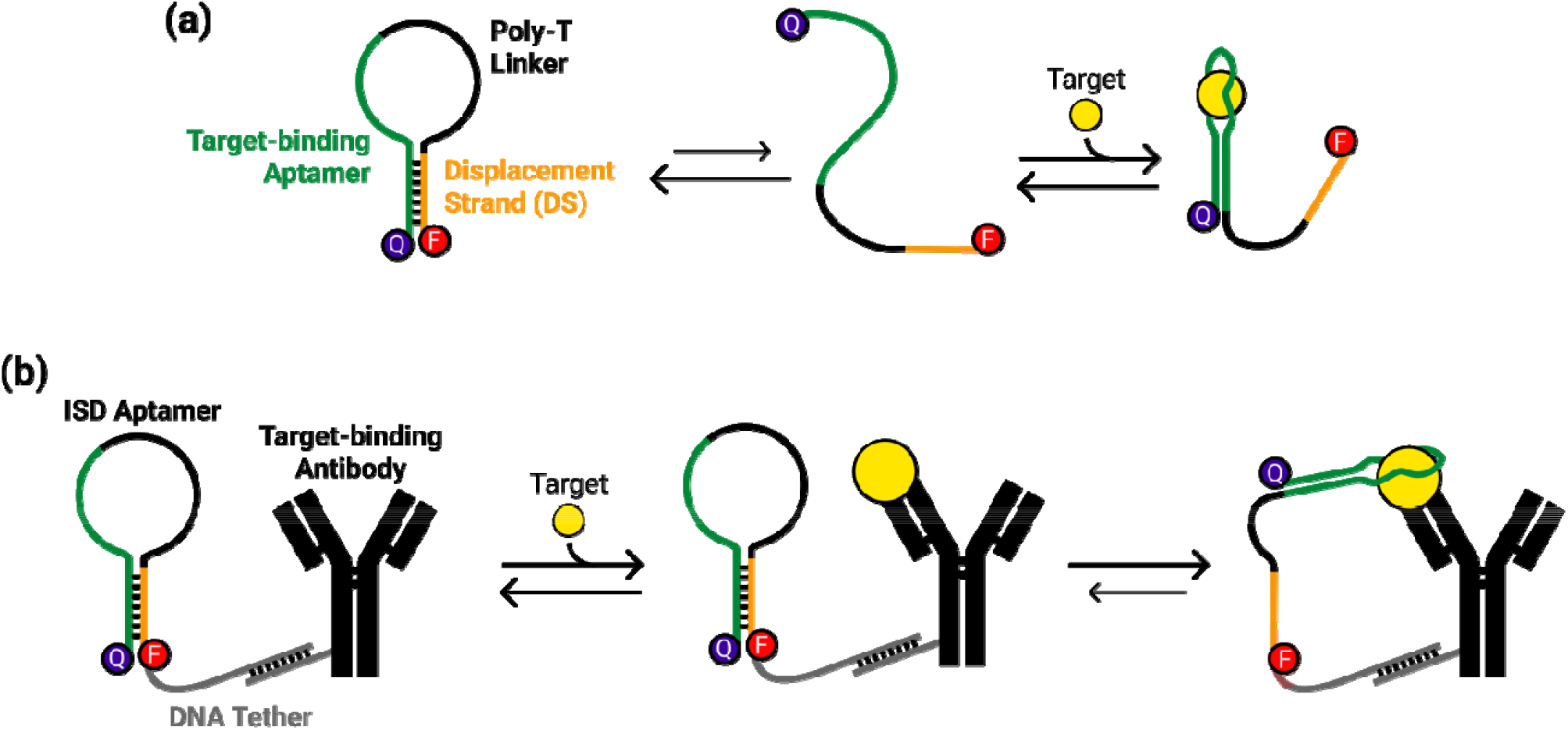
(a) Schematic of the intramolecular strand displacement (ISD) aptamer switch mechanism. In the ISD switch, an aptamer is linked to a partially complementary displacement strand (DS) via a poly-T linker, and the two ends of the DNA molecule are labeled with a fluorophore and quencher moiety. Target binding leads to release of the DS and physical separation of these two elements, resulting in increased fluorescence. (b) Schematic of the Programmable Antibody and DNA Aptamer Switch (PANDAS) mechanism. In the PANDAS design, the ISD aptamer is coupled to the Fc domain of an antibody by a 30-nt oligonucleotide tether. The antibody will generally have much higher affinity than the aptamer, capturing the free target first and bringing it into close proximity with the ISD, allowing the captured target molecule to bind the ISD and induce a conformational change that produces a fluorescent readout.

The antibody component is site-specifically modified with a 30-nt DNA tether at its Fc domain, which is in turn tethered to the aptamer switch component via hybridization to a complementary tether sequence that is appended to the fluorophore-tagged DS end of the ISD aptamer (all sequences shown in **Supplementary Table S1**). This site-specific tethering ensures reproducible interactions between the target-binding sites of the aptamer and antibody, thereby enabling cooperative binding and reliable aptamer signaling. The modularity of this design allows for the ready substitution of different aptamer switches or antibody candidates within the PANDAS construct, enabling rapid testing of different switch configurations.

### Thermodynamic modeling of the PANDAS binding mechanism

To understand the degree of aptamer affinity enhancement that could be achieved, we developed a thermodynamic model of target-induced binding and switching for the PANDAS construct. We modeled the construct as two linked receptors: the aptamer and the antibody. The binding and switching of the chimeric construct are modeled in terms of target-dependent equilibria between five states, which in turn represent different combinations of target binding for the two receptors (**Fig. 2a**). In the majority of cases, the antibody affinity will be higher than that of the ISD aptamer, such that the antibody will capture the target first. The effective concentration of target in the proximity of the ISD aptamer will be dramatically increased due to the aptamer-antibody linkage, and this high effective concentration will trigger aptamer binding and fluorescent signaling. We assume that the switch produces fluorescent signal only in states where the aptamer is bound to the target. In this model, the function of the PANDAS construct is dictated by three properties of the switch:, and is the effective concentration of target experienced by the aptamer when the target is already bound to th antibody, while *K*_*DAp*_ and *K*_*DAb*_ are the equilibrium dissociation constants of the aptamer and antibody, respectively. While and are not easily tunable within the PANDAS design, the abovementioned ISD tuning strategy allows us to control.

**Figure 2.**
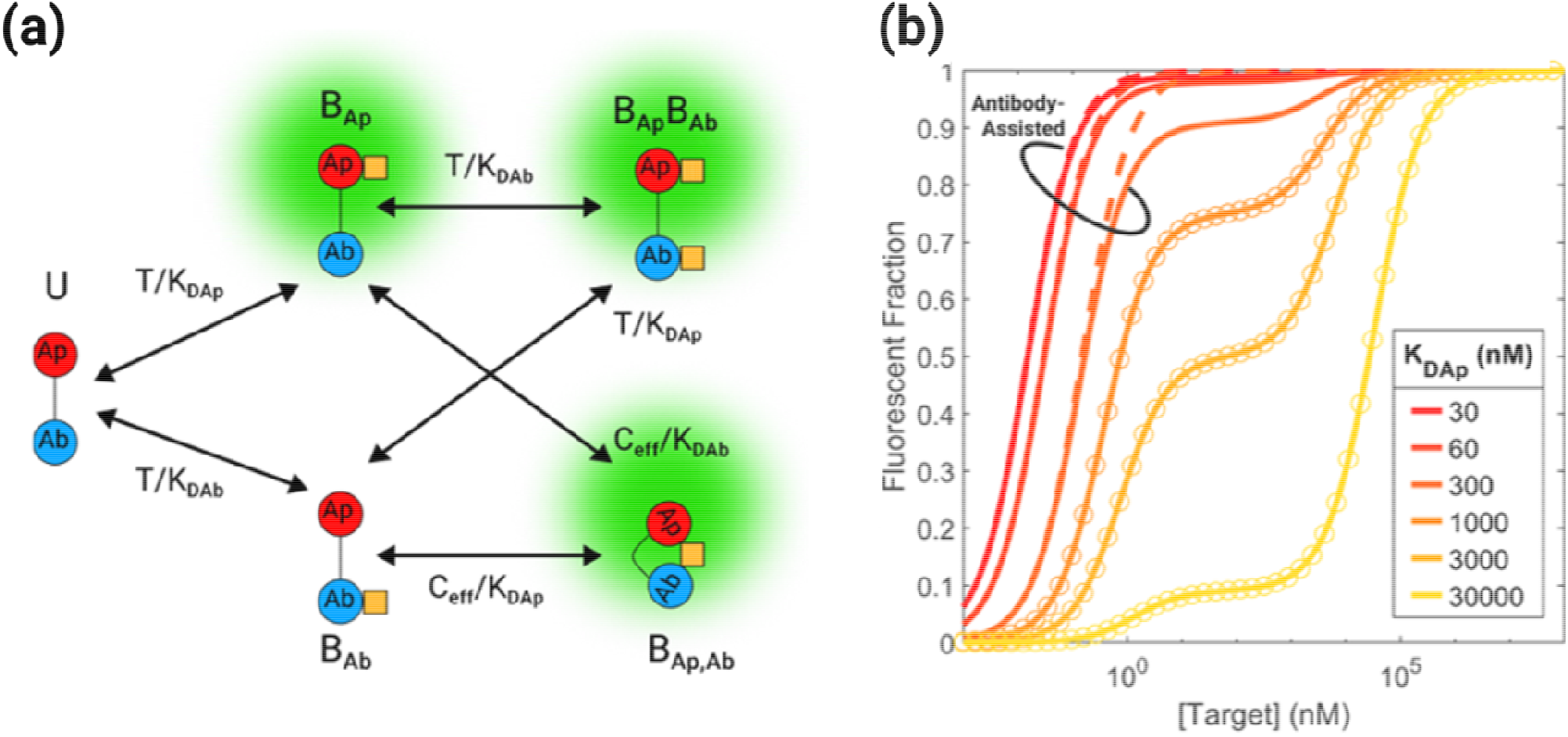
Modeling the thermodynamic binding behavior of the PANDAS construct. (a) Our simplified PANDAS model considers five conformations for the aptamer (Ap), antibody (Ab), and target (yellow square): unbound (U), target singly bound to the aptamer (B_Ap_) or antibody (B_Ab_), the bivalently bound conformation (B_Ap,Ab_), and the doubly bound conformation (B_Ap_B_Ab_). (b) Tuning of PANDAS affinity by changing ISD aptamer affinity. The plot shows the fraction of fluorescently active constructs at various concentrations of target in a scenario where K_DAb_ = 3 nM, C_eff_ = 3 μM, and K_DAp_ ranges from 30 nM–30 μM. Solid lines represent plots of the full thermodynamic binding model. Dashed lines represent the simplified single-isotherm approximation used when *K*_*DAp*_ ≪ *C*_*eff*_. ⨀ markers represent the dual-isotherm approximation used when *K*_*DAp*_ ≥ *C*_*eff*_.

We leveraged this model to predict the binding response of the PANDAS construct for different values of *K*_*DAp*_ (**Fig. 2b**; see **SI, Appendix 1** for details). We identified two different binding regimes for the PANDAS construct. The ‘antibody-assisted’ regime corresponds to values of *K*_*DAp*_ where *C*_*eff*_ ≫ *K*_*DAp*_ ≫ *K*_*DAb*_. In this case, once the target molecule is bound to the antibody, the aptamer readily binds it as well. As such, the aptamer-bound-only state, B_Ap_, does not contribute meaningfully to the binding signal, and the fraction of fluorescent constructs is well-approximated by a single isotherm:

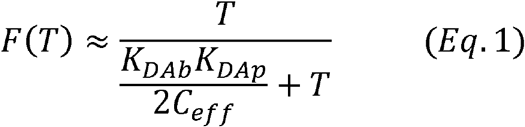

This approximation is shown as dashed lines in **Figure 2b**. In the antibody-assisted regime, the PANDAS effective affinity (*K*_*DEff*_) is given by *K*_*DAb*_ *K*_*DAp*_/(2*C*_*eff*_), and thus depends linearly on the affinity of the ISD aptamer component, scaled by an enhancement factor of *K*_*DAb*_/(2*C*_*eff*_). As *K*_*DAp*_ approaches or exceeds *C*_*eff*_ (*K*_*DAp*_ ≥ *C*_*eff*_ ≫ *K*_*DAb*_), the B_Ap_ state becomes more prominent, as not all instances of target binding to the aptamer are necessarily triggered by an initial antibody binding event. In this ‘antibody-independent’ regime, the fraction of fluorescent PANDAS constructs is better approximated by a dual-isotherm given by:

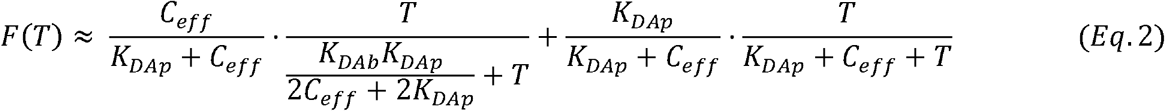

This approximation is shown in **Figure 2b** as ⊙ symbols. The first, high-affinity isotherm describes the antibody-assisted signal, while the latter, low-affinity isotherm describes the contribution of the B_Ap_ state. When *K*_*DAp*_ ≫ (*i*.*e*., aptamer affinity is so low that the proximity of antibody-bound targets is insufficient to trigger aptamer binding), the PANDAS binding response depends only on the affinity of the aptamer. As such, aptamers with extremely low affinities (*e*.*g*., in the mM range) will not experience sensitivity enhancement from a tethered high-affinity antibody, as the local enhancement of target concentration from proximity will be insufficient to trigger aptamer binding. Fortunately, most protein-binding aptamers exhibit affinities in the sub-micromolar range and are thus suitable for use in the PANDAS design.

### Testing the PANDAS construct with a thrombin model system

As a proof-of-concept demonstration of the affinity enhancement that can be achieved with PANDAS, we designed a fluorescent sensor for thrombin, an important factor in the blood clotting process for which several well-characterized aptamers and monoclonal antibodies have been described^20–25^. Using flow cytometry, we performed pairwise screening of potential aptamer (TBA, HD22) and antibody candidates (5020, F1) to determine an appropriate aptamer-antibody pair and determined that the combination of the 5020 antibody and TBA aptamer achieved the most robust sandwich binding of thrombin (**Supplementary Fig. S2, S3**).

We then converted the TBA sequence into an ISD-based fluorescent molecular switch. Initially, we selected a DS length of 7 nt to achieve a high amplitude switching signal with moderate binding affinity. Since the PANDAS enhancement mechanism depends on the affinity difference between the aptamer and antibody, we performed biolayer interferometry (BLI) experiments to determine the affinities of our individual switch components (**Supplementary Fig. S4–S6**). As expected, our TBA-based ISD aptamer switch exhibited lower affinity (K_D_ = 246 nM) compared to the parent aptamer (K_D_ = 114 nM), whereas the 5020 antibody had a much higher affinity (K_D_ = 2.76 nM) than the aptamer switch. These results supported the suitability of this affinity reagent pair to achieve cooperative enhancement of aptamer binding at low target concentrations based on our model. We then site-specifically modified 5020 antibodies at their Fc region with a 30-nt tether oligo at a 1:1 ratio (**Supplementary Fig. S7**). Subsequently, the TBA ISD aptamer sequence was hybridized to this tether oligo to produce the complete PANDAS construct. The sequences were designed such that the tether between aptamer and antibody consisted of a 30-nt hybridization region as well as a 5-nt single stranded poly-T spacer, such that the overall 35-nt tether region is sufficiently long to achieve robust antibody-aptamer hybridization with minimal dissociation, while not being so long that steric interference might impede sandwich binding. The assembled PANDAS construct was then immobilized onto microbeads coated with protein G, which binds to the antibody Fc region. These PANDAS-conjugated beads were incubated for 30 min in buffer spiked with a range of thrombin concentrations, followed by measurement of fluorescent switching using a flow cytometer. As a control, we generated a non-cooperative PANDAS construct in which the TBA ISD was coupled to an anti-TNFL □ antibody that does not bind the thrombin target and challenged this construct with the same range of thrombin concentrations. This assay showed that our PANDAS construct had a K_D_ of ∼1.7 nM (**Fig. 3a**), achieving nearly 100-fold greater sensitivity relative to the non-cooperative control construct (K_D_ ∼139.3 nM). We also ruled out effects of non-specific protein binding by testing the thrombin antibody-based PANDAS construct against bovine serum albumin (BSA), as well as effects of antibody binding without specific aptamer binding by challenging the anti-TNFL □-based PANDAS construct with various concentrations of TNFL □, and in both cases we observed no fluorescent signal. These experiments confirmed that signal generation was dependent on aptamer recognition of the target, whereas the affinity and sensitivity increase was contingent upon the specific and cooperative binding of both antibody and aptamer.

**Figure 3.**
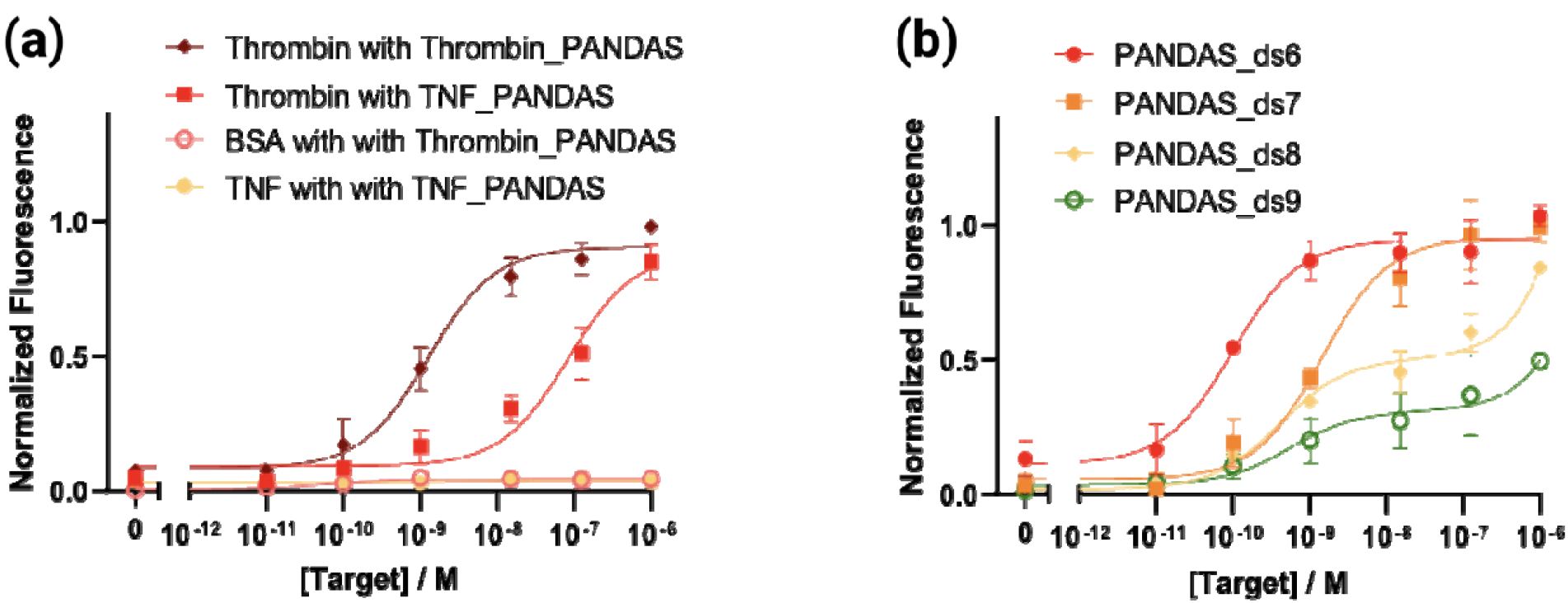
Specificity and tunability of PANDAS constructs. (a) Testing PANDAS specificity and the impact of cooperative binding. We used a flow-cytometry-based assay to measure the binding-induced fluorescent readout from our thrombin PANDAS in the presence of thrombin and bovine serum albumin (BSA) compared to a non-cooperative PANDAS construct with a mismatched anti-TNF-α antibody exposed to thrombin or TNF-α. (b) Tuning the binding affinity of the thrombin PANDAS construct. We tailored the length of the DS sequence in our PANDAS to range from 6–9 nt and then used flow cytometry to measure fluorescence after 30 min of incubation with varying concentrations of thrombin. All datapoints and error bars represent the mean and standard deviation of three replicates.

### Rational sequence-based tuning of the PANDAS switch

Building off insights from our thermodynamic model, we developed a range of PANDAS switches in which we tuned the ISD design by extending or shortening the DS. We predicted that elongating the DS would stabilize its hybridization to the aptamer and increase its competition with target binding, thereby decreasing the thrombin affinity of both the aptamer and the overall PANDAS construct. Conversely, shortening the DS should weaken intramolecular competition, leading to higher thrombin affinity for the aptamer and PANDAS. We selected DS lengths ranging from 6–9 nt (using the naming convention Aptamer_ds#), immobilized these constructs onto beads, and quantified their thrombin-binding affinity using the flow cytometry-based approach described above.

As expected, decreasing the DS length increased ISD aptamer binding affinity and thus shifted the dynamic range towards more accurate sensing of lower target concentrations (**Fig. S8**). For example, decreasing the DS length from 7 to 6 nt led to a 15-fold decrease in *K*_*D*_ from 139.3 nM to 9.01 nM. Conversely, extending the DS length from 7 to 9 nt weakened the aptamer’s sensitivity considerably, yielding binding curves for which it proved difficult to achieve saturation. We next assembled these ISD constructs into full PANDAS constructs with the anti-thrombin antibody, and again assessed their binding via flow cytometry (**Fig. 3b**). Between PANDAS_ds6 and PANDAS_ds9, we observed a range of effective affinities spanning from 0.1–1,000 nM. These experimentally derived binding curves aligned quite well with the two binding regimes predicted by our model. PANDAS_ds6 and PANDAS_ds7 approximated single-isotherm binding, with increased sensitivity as a result of antibody-assisted binding, yielding *K*_*D*_ values of 97.72 pM and 1.43 nM, respectively. In contrast, PANDAS_ds8 and PANDAS_ds9 exhibited two-stage binding with a broader dynamic range due to antibody-independent binding. The ability to rationally tune this response from high sensitivity to wide dynamic range sensing could prove valuable for the optimization of PANDAS constructs for specific biomolecular detection applications. Although PANDAS_ds6 exhibited the best sensitivity, its background was also high due to the reduced stability of DS hybridization. For the remainder of this work, we chose PANDAS_ds7 for further characterization because of its low background, high signal gain, and relatively high thrombin sensitivity.

To evaluate the kinetic response of this construct, we dispensed PANDAS_ds7-coupled beads into a 1 μM thrombin-spiked buffer and incubated them for three minutes to allow them to equilibrate. We then washed the beads with thrombin-free buffer and periodically interrogated the PANDAS fluorescent binding response via flow cytometry to observe the target dissociation signal (**Fig. 4a**). As a control, we repeated this experiment by washing the beads with the same 1 μM thrombin-containing solution, which should result in minimal dissociation. We measured a dissociation rate of k_off_ = 8.3 × 10^−4^ s^-1^, which indicates that the PANDAS construct maintains a reasonable off-rate that is suitable for tracking endogenous target concentration changes in continuous sensing applications, given an antibody with typical binding kinetics. We also evaluated the reversibility of binding by deploying the PANDAS-coupled beads into alternating solutions of 1 μM thrombin and buffer, incubating for 15 minutes each time, and then interrogating the construct via flow cytometry (**Fig. 4b**). The PANDAS receptor was able to track these changes in concentration over many cycles, with the binding signal returning to near-baseline levels every time.

**Figure 4.**
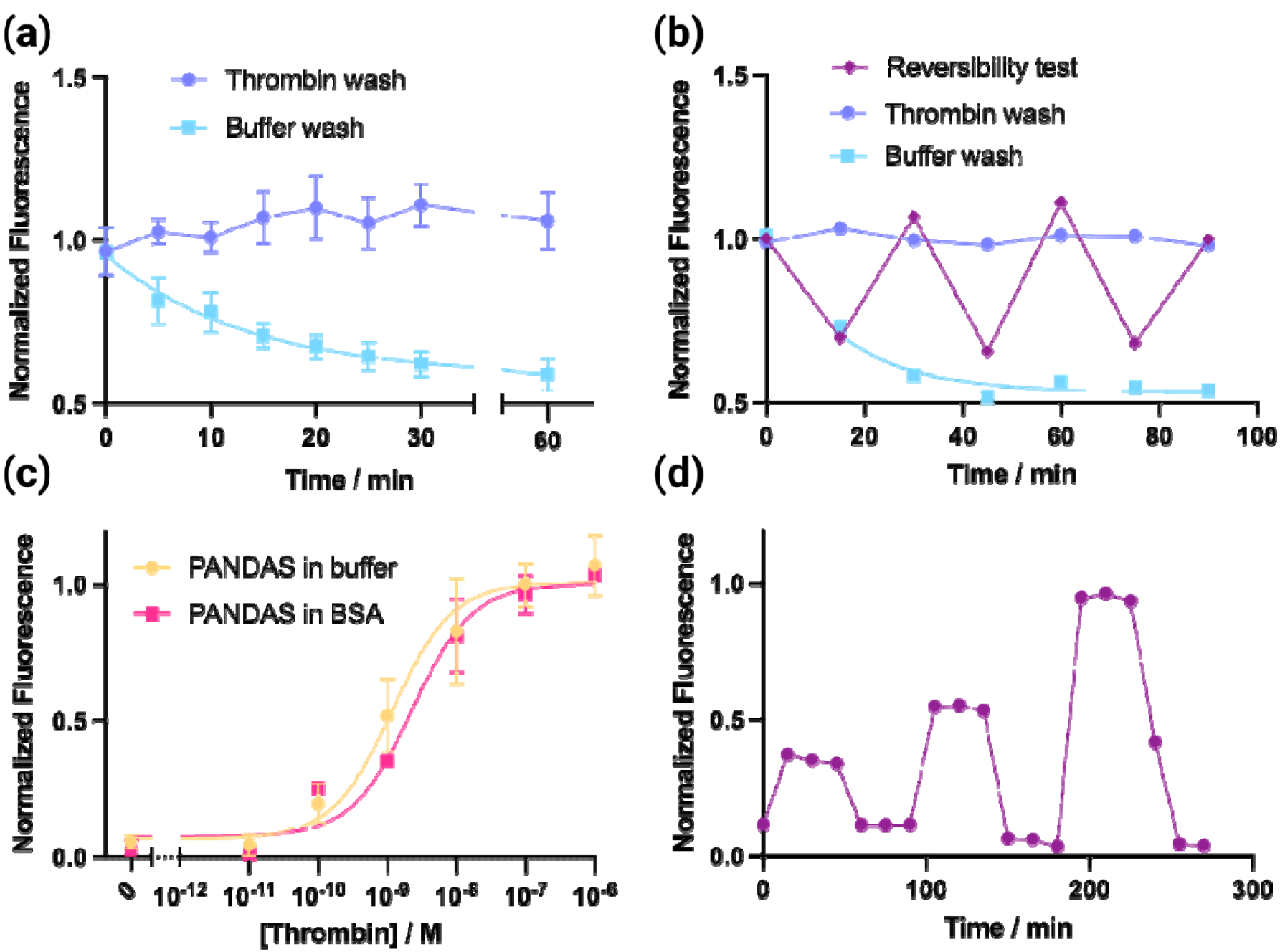
PANDAS kinetics and reversibility of target binding. (a) To measure the kinetics of PANDAS_ds7, we generated dissociation curves by washing thrombin-saturated PANDAS beads with buffer or 1 μM thrombin and measuring fluorescence every three minutes. (b) To assess reversibility, we transferred PANDAS_ds7-coupled beads between buffer and 1 μM thrombin, collecting fluorescence measurements 15 minutes after each shift. As controls, we collected fluorescence measurements from PANDAS-ds7 beads that were maintained in either buffer or 1 μM thrombin. (c) PANDAS deployment in an interferent-rich environment. PANDAS_ds7 maintained consistent thrombin-binding performance in buffer alone or containing 530 μM albumin. (d) We also tested the sensor’s capacity for continuous sensing of a physiological range of thrombin concentrations in buffer containing 530 μM albumin, with data collected every 15 min. Figure a and c datapoints and error bars respectively represent the mean and standard deviation of three replicates.

Finally, we assessed the PANDAS construct’s performance as a continuous sensor in complex samples. Working with the protein-rich matrix of blood, serum, or plasma would have required a careful selection of samples to avoid the influence of proteins that interfere with our ability to accurately measure spiked-in thrombin concentrations. Endogenous antithrombin III (ATIII), heparin cofactor II (HCII), and alpha-2-macroglobulin (A2M) can all form complexes with thrombin and thereby interfere with the function of our PANDAS sensor^26–28^. We therefore used a surrogate protein-rich biofluid of defined content, challenging our switch with buffer containing physiologically representative concentrations of BSA (∼530 μM) as a surrogate for human albumin—a large protein that is prone to nonspecific binding and composes 60% of total plasma protein content^29^. We measured the fluorescent response of PANDAS_ds7 in this albumin-rich solution using the same bead-based fluorescent assay described above, with 30-min sample incubation time. The response to thrombin in the presence of albumin did not differ significantly from its performance in buffer (**Fig. 4c**), indicating that the PANDAS design can leverage the antibody’s excellent specificity to facilitate sensitive aptamer-based detection in complex samples. This is especially notable as the TBA aptamer has been reported to cross-react with BSA with micromolar affinity^30^. Finally, to test our construct’s capacity for time-resolved continuous sensing in complex samples, we sequentially incubated our beads with varying physiological-range concentrations of thrombin in the presence of 530 μM albumin and measured the switch response every 15 min (**Fig. 4d**). We observed stable concentration-dependent measurements over the entire 270-minute experiment, demonstrating the potential for continuous monitoring of analyte concentrations with our PANDAS receptor.

## Conclusions

Molecular switches are valuable tools for the rapid detection and continuous monitoring of diverse analytes, but existing approaches for generating such reagents suffer from notable limitations. Aptamer-based switches such as the ISD constructs used in this work are relatively straightforward to engineer from existing aptamers, but the engineering process tends to sacrifice the sensitivity of the final molecular switch. In contrast, antibodies generally offer excellent affinity and specificity in the context of endpoint-based measurements, even in complex samples, but it remains challenging to achieve continuous analyte detection using antibody-based assays beyond a few preliminary demonstrations^10^. Our PANDAS design offers the best of both worlds—greatly enhancing the affinity and specificity of an ISD-based aptamer switch by linking it to a target-specific antibody. In this design, the antibody enables an initial high-affinity binding event, and the resulting increased local concentration then favors cooperative binding by the aptamer component. The aptamer produces a fluorescent readout for sensing and can readily be tuned by modulating the ISD design to optimize its sensitivity. Using thrombin as a model analyte, we demonstrated that the PANDAS design can enhance the sensitivity of protein detection by ∼100-fold relative to the ISD-based aptamer switch on its own. The resulting PANDAS are highly analyte-specific, leveraging two-site binding from both elements to recognize their target with high sensitivity. We show that PANDAS switches can achieve rapid and reversible switching over multiple cycles of target binding and washing, suggesting the feasibility of applying these receptors towards continuous detection with minute-scale temporal resolution. We also demonstrate the ability to monitor fluctuating levels of thrombin within a physiological concentration range in complex samples containing high levels of interferent protein. Critically, the PANDAS design should be compatible with aptamer switches developed through a wide range of approaches, and thus offers the potential to enhance the sensitivity of sensors for a variety of lower-abundance biomarkers.

It should be noted that this is only an initial demonstration of the PANDAS concept, and additional work will be needed to maximize the utility of this reagent design. As the design is extended to more protein biomarkers, aptamer function must be carefully optimized to ensure both high sensitivity and suitable temporal resolution of measurements. To this end, the properties of the PANDAS thrombin sensor and previous demonstrations of ISD aptamer tunability indicate that it should be feasible to balance sub-nanomolar temporal resolution with minute-scale kinetic responses by tuning the DS and linker components within the aptamer^17^. While we have demonstrated the reversibility and minute-scale measurement kinetics of the PANDAS target-binding response, achieving continuous real-time sensing will require a more sophisticated sensor design that enables surface-based fluorescent measurements, such as the optical fiber sensors recently demonstrated by our group^31^. Such surface-based sensors will also enable further investigation of the quantitative performance of PANDAS-based measurements in more complex biological specimens. Although we were able to show that sensing performance was unimpeded by high background levels of interferent protein, it remains to be seen whether this system will remain equally robust in the context of blood and other complex biological matrices that are incompatible with bead-based assays. Overall, we see considerable future potential for this sensor design approach, which we believe should be broadly generalizable across a range of molecular targets and offers the opportunity to convert moderate-quality aptamers into highly useful reagents for achieving sensitive molecular detection in interferent-rich specimens.

## Supporting information

Supplementary Figures, Tables, and Discussion

## Acknowledgements

HTS gratefully acknowledges financial supported by Helmsley Charitable Trust, the Stanford Maternal and Child Health Research Institute (MCHRI) and the Wellcome LEAP SAVE program. I.A.P.T. was supported by the Medtronic Foundation Stanford Graduate Fellowship and the Natural Sciences and Engineering Research Council of Canada (NSERC, 416353855).

