## Supplementary Figures, Tables, and Discussion for "Aptamer-antibody chimera sensors for sensitive, rapid and reversible molecular detection in complex samples"

### Materials and Methods

#### Reagents and materials

DNA oligonucleotides with fluorophore-quencher modifications were purchased from Integrated DNA Technologies. All sequences are shown in **Table S1**. Unless otherwise specified, all other reagents were ordered from Thermo Fisher Scientific. Oligonucleotide concentrations were determined by UV spectroscopy using a Nanodrop ND2000 spectrophotometer. Thrombin (HCT-0020 human α-thrombin) was purchased from Haematologic Technologies. Thrombin monoclonal antibody 5020 (Catalog # MA1-43019) was purchased from Thermo Fisher Scientific. Thrombin monoclonal antibody F1 (sc-271449) was purchased from Santa Cruz Biotechnology. TNF-α antibody was purchased from Biolegend (Catalog # 502802). Molecular biology grade BSA was purchased from New England Biolabs. Protein G Dynabeads (Catalog # 10-004-D) were purchased from Invitrogen. PBS, Salmon Sperm DNA, goat IgG, EDTA, NuPAGE sample reducing agent, NuPAGE LDS Sample Buffer (4x), NuPAGE 4 to 12% Bis-Tris mini protein gels, 20x NuPAGE MES SDS running buffer, SeeBlue pre-stained protein standard, SimplyBlue SafeStain, UltraPure TBE buffer (10X), Novex TBE-Urea Gels (10%), GelStar nucleic acid gel stain, and Tween 20 were purchased from Thermo Fisher Scientific.

#### Screening aptamer-antibody pairs for the PANDAS construct

We tested two Cy5-labeled thrombin aptamer sequences (TBA and HD22) and two different anti-thrombin antibodies (5020 and F1). 200 nM antibody was incubated with 10 µL protein G Dynabeads in 50 µL 1× ISD binding buffer (10 mM Tris–HCl, pH 7.5 and 6 mM $MgCl_2$) for 30 min at room temperature. The beads were washed three times to remove any nonspecific binding, and then incubated with 450 nM aptamer in the presence or absence of 300 nM thrombin in a final volume of 50 µL ISD buffer for 30 min at room temperature.

#### PANDAS assembly

We used the SiteClick antibody azido modification kit (S20026, Thermo Fisher Scientific) to modify the antibody Fc region in a site-specific fashion. 100 µg of antibody was treated with 10 µL β-galactosidase in 20 µL 1x PBS buffer at 37 °C overnight to remove the terminal β-Dgalactoside. The uncapped antibodies were then treated with 80 µL GalT transferase and UDP-GalNAz in 1x Tris buffer at 30 °C overnight to add azide caps. 50K Amicon purification columns were used to remove excessive UDP-GalNAz. 50 µL of 100 µM amine-modified tether oligonucleotides (Ab2-tether) were treated with 100 µL 17 mM DBCO-NHS at room temperature overnight to equip these oligos with DBCO moieties. The oligos were then ethanol precipitated to remove excess reactants and resuspended in 1× ISD buffer. 10 µL of 100 µM DBCO-modified tether oligonucleotides were reacted with 10 µL of 10 µM azido-antibody in 1× PBS buffer, and the degree of antibody labeling was determined by running a reducing NuPAGE gel (see below). The antibody-tether complexes were stored at 4 °C until further use. To make fresh PANDAS, 2 µL of 5 µM antibody-tether complexes was hybridized to 1 µL of 100 µM aptamers in hybridization buffer (1x PBS, 1 mg/mL BSA, 0.05% Tween 20, 15 µg/m goat IgG, 0.1 mg/mL salmon sperm DNA, and 5 mM EDTA) overnight. 10 pmol of PANDAS chimera was then captured by 10 µL of prewashed protein G Dynabeads and washed three times with ISD buffer to remove excess aptamers.

**Antibody labeling quantification in reducing gels**

We quantified the degree of antibody oligo labeling with a 10% NuPAGE gel. 1 µL NuPAGE reducing reagent was incubated with 12 pmol unmodified antibody or click-modified antibody in a total volume of 10 µL 1X NuPAGE sample loading buffer at 95 °C for 5 min. The samples were then quickly loaded onto the 10% Bis-Tris mini protein gel and run at 200 V for 30 min. After 30 min, the gel was transferred to 100 mL SimplyBlue SafeStain staining solution for another 30 min. The gel was rinsed with water twice to destain before imaging on a ChemiDoc MP System (Bio-Rad Laboratories).

**Biolayer interferometry (BLI) analysis of PANDAS components**

BLI affinity measurements were done on an Octet Red 96 instrument (Pall ForteBio), with streptavidin sensors (SA, Sartorius) and Fc capture sensors (AMC, Sartorius). Sensors were pre-hydrated in ISD buffer for at least 20 min before use. 200 µL of 20 nM biotinylated aptamers or antibodies in ISD buffer were used for ligand loading to reach ~0.5 nm loading levels. After ligand loading, sensors were dipped in ISD binding buffer containing various thrombin concentrations

(25, 50, 100, or 200 nM for aptamer measurements; 10, 20, 40, or 80 nM for antibody measurements) for 300 sec and then dipped in ISD buffer for 600 sec. To avoid nonspecific binding, every thrombin concentration was measured with both assay and reference sensors. Final signals were processed via signal subtraction from reference sensors.

**Measurements of effective binding affinity**

To obtain binding curves, 50 μL reactions were prepared in 1× ISD binding buffer with 10 nM PANDAS-coupled beads and thrombin concentrations in the range of 1 nM–1 μM for 30 min at room temperature. For ISD-only tuning experiments, we used a TNF-α antibody that was shown have no affinity for thrombin as a dummy antibody in the final PANDAS construct. For baseline determination with the PANDAS alone, 50 μL of 10 nM PANDAS-coupled beads were prepared in ISD binding buffer without thrombin. The mean fluorescent intensities for all samples were measured at 25 °C on a flow cytometer (BD Accuri C6 Plus). All samples were prepared and analyzed in triplicate.

For the reversibility experiments, three samples of 10 nM PANDAS-coupled beads and 1 μM thrombin in 50 μL of 1× ISD buffer were incubated for 30 min at room temperature to reach saturation. The sample tubes were then put in a magnetic rack (Life Technologies) for 2 min to separate PANDAS beads and sample liquid. For the first sample, we maintained saturation with 50 μL of 1 μM thrombin in ISD buffer added to the beads every 15 min. The second sample was for reversibility testing, and 50 μL of ISD buffer or 1 μM thrombin in ISD buffer were added every 15 min in an alternating fashion. The third sample was used to measure dissociation only, with 50 μL of 1× ISD binding added to the PANDAS beads every 15 min. The mean fluorescent intensities for all samples were measured at 25 °C on a flow cytometer.

**Signal normalization for thermodynamic plots**

We applied single-site hyperbolic binding model to fit and normalize ISD data and PANDAS_ds6 and _ds7 data from the flow cytometer for ease of comparison:

$$y = (B_{max} - y_0)\frac{x}{x + K_D^{eff}} + y_0$$

We applied a two-site binding model to fit and normalize PANDAS_ds8 and _9 data from the flow cytometer:

$$y = (B_{max}Hi - y_0)\frac{x}{x + K_D^{Hi}} + (B_{max}Lo - y_0)\frac{x}{x + K_D^{Lo}} + y_0$$

The average fluorescence value from three replicates was divided by $B_{max}$ before plotting. Error bars were calculated as propagation of errors for standard deviation from the average fluorescence and error in the fit for $B_{max}$.

### Supplementary Table

**Table S1. Oligo sequences**

| Name | Sequence |
|---|---|
| Ab2 tether | /5AmMC12/TCAACAATAGATAAGTCCTGAACAAGAAAA |
| ThrombinISD_5T_Tether_Cy5_DS6 | /5IAbRQ/TCACTGTGGTTGGTGTGGTTGGTTTTTTTTTTTTACAGTG/iCy5/TTTTTTTTTCTTGTTCAGGACTTATCTATTGTTGA |
| ThrombinISD_5T_Tether_Cy5_DS7 | /5IAbRQ/TCACTGTGGTTGGTGTGGTTGGTTTTTTTTTTTTCACAGTG/iCy5/TTTTTTTTTCTTGTTCAGGACTTATCTATTGTTGA |
| ThrombinISD_5T_Tether_Cy5_DS8 | /5IAbRQ/TCACTGTGGTTGGTGTGGTTGGTTTTTTTTTTTTCCACAGTG/iCy5/TTTTTTTTTCTTGTTCAGGACTTATCTATTGTTGA |
| ThrombinISD_5T_Tether_Cy5_DS9 | /5IAbRQ/TCACTGTGGTTGGTGTGGTTGGTTTTTTTTTTTTACCACAGTG/iCy5/TTTTTTTTTCTTGTTCAGGACTTATCTATTGTTGA |
| TBA-Bt | /5BiotinTEG/GGTTGGTGTGGTTGG |
| ISD-Bt | /5BiotinTEG/TCACTGTGGTTGGTGTGGTTGGTTTTTTTTTTTTACAGTG |
| TBA-Cy5 | /5Cy5/GGTTGGTGTGGTTGG |
| HD22-Cy5 | /5Cy5/AGTCCGTGGTAGGGCAGGTTGGGGTGACT |
| **Oligo Modifications** | |
| /5AmMC6/ | 5' amino modifier C6 from IDT |
| /5IAbRQ/ | 5' Iowa Black® RQ Quencher from IDT |
| /iCy5/ | Internal Cy5 from IDT |

| | |
|---|---|
| /5BiotinTEG/ | 5’ Biotin-TEG from IDT |
| /5Cy5/ | 5’ Cy5 from IDT |

#### Supplementary Figures

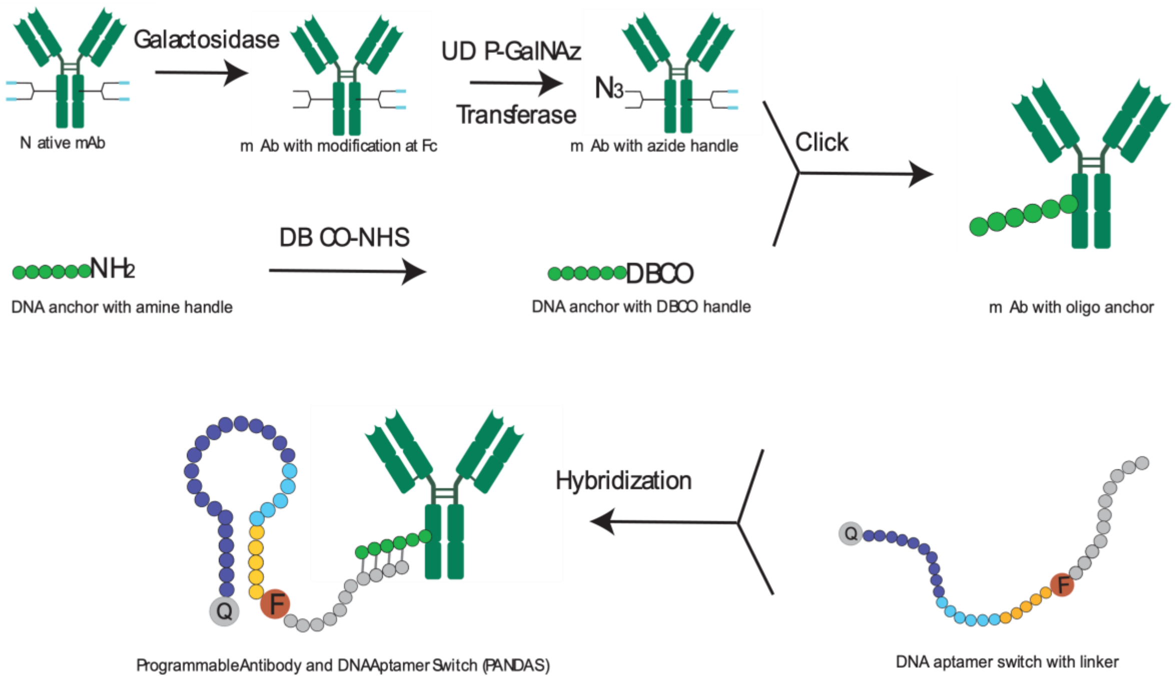


**Figure S1**. Assembly process for the PANDAS construct. An azide handle is attached to the Fc domain of a native antibody by performing galactosidase digestion followed by UDP-GalNAz addition. In parallel, a DBCO handle is attached to an amine-modified DNA tether ('Ab2 tether' in **Table S1**) using DBCO-NHS. This DNA tether is then conjugated to the antibody's azide group via a copper-free click reaction. This oligonucleotide-modified antibody can now be hybridized to the aptamer switch, yielding the final PANDAS construct.

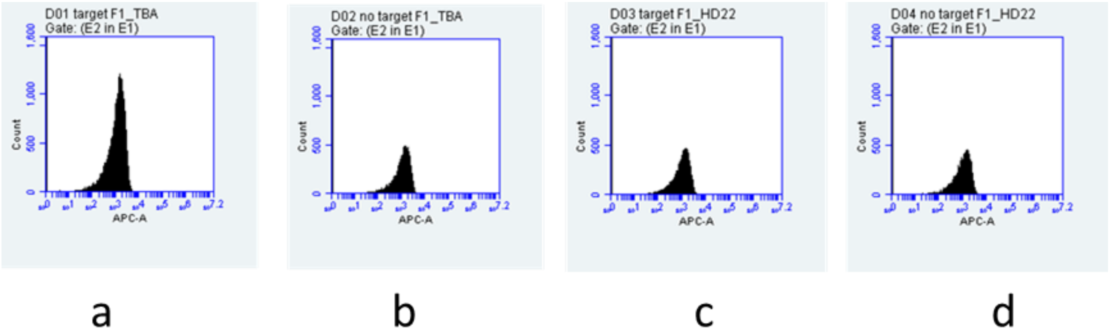


**Figure S2**. Assessing PANDAS antibody-aptamer compatibility. 200 nM antibody F1 was immobilized to protein G beads, and then incubated with 450 nM aptamer TBA (**a, b**) or HD22 (**c**, **d**) in ISD buffer with (**a**, **c**) or without (**b**, **d**) 300 nM thrombin. Each reaction was then analyzed via flow cytometry.

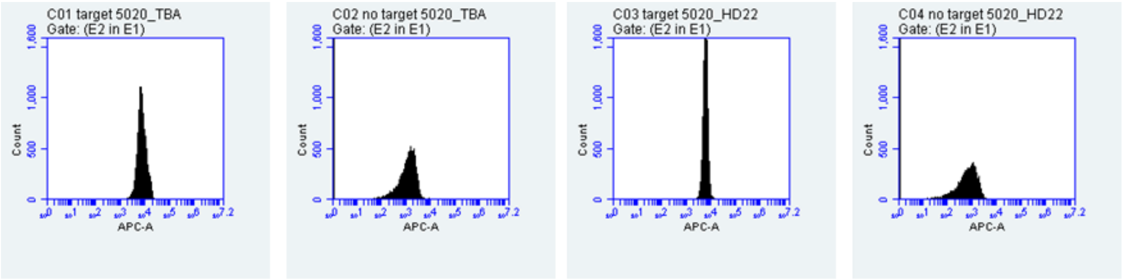


a b c d

**Figure S3**. Assessing PANDAS antibody-aptamer compatibility. 200 nM antibody 5020 was immobilized to protein G beads, and then incubated with 450 nM aptamer TBA (**a, b**) or HD22 (**c**, **d**) in ISD buffer with (**a**, **c**) or without (**b**, **d**) 300 nM thrombin. Each reaction was then analyzed via flow cytometry.

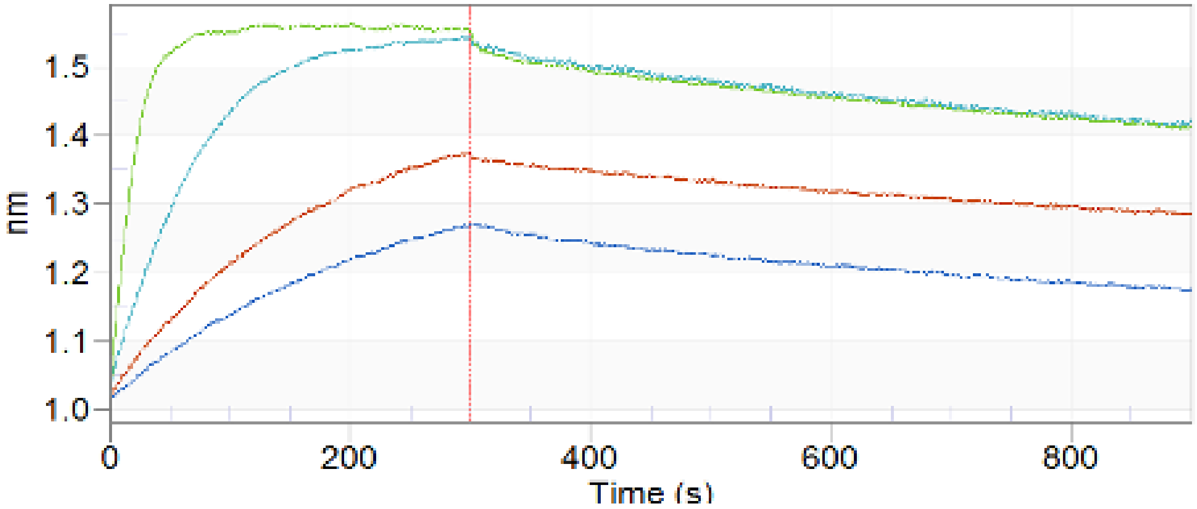


**Figure S4**. BLI affinity measurements of antibody 5020. AMC tips were pre-hydrated in ISD buffer for 20 minutes and then in 20 nM 5020 antibody to achieve 0.5 nm loading level. After ligand loading, sensors were dipped in 10, 20, 40, or 80 nM thrombin for 300 sec and then dipped in buffer for 600 sec. Signals were normalized by subtraction of the signal from reference tips. ForteBio Octet Data Analysis software was used to align and fit the data in 1:1 binding mode ($K_{on}$ = 6.09 x $10^5$ $M^{-1}s^{-1}$, $K_{off}$ =1.68 x $10^{-3}$ $s^{-1}$ and $K_D$=2.76 x $10^{-9}$ M).

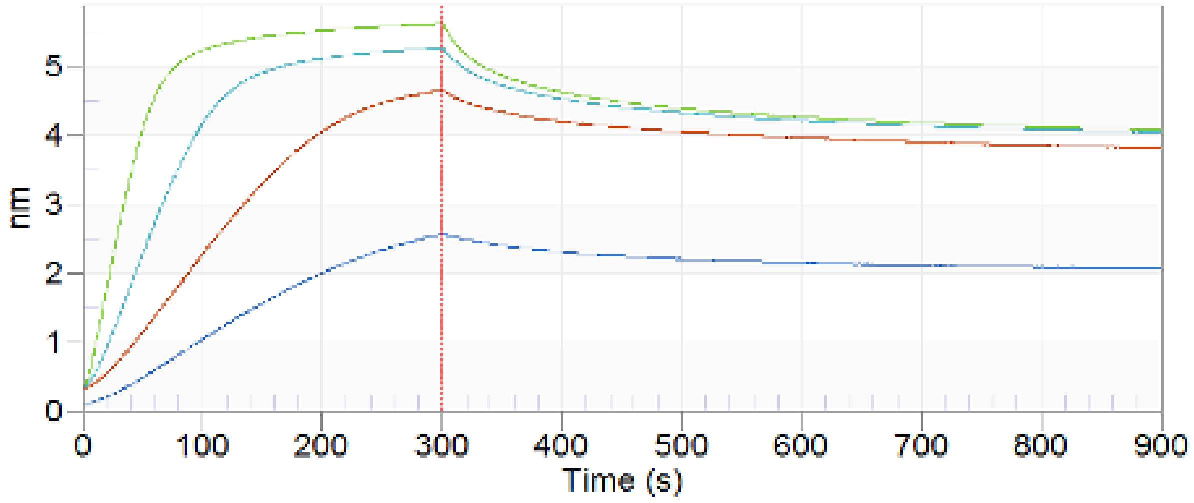


**Figure S5**. BLI affinity measurements of aptamer TBA. SA tips were pre-hydrated in ISD buffer for 20 minutes and then in 20 nM TBA to achieve 0.5 nm loading level. After ligand loading, sensors were dipped in 25, 50, 100, or 200 nM thrombin for 300 sec and then dipped in buffer for 600 sec. Signals were normalized by subtraction of the signal from reference tips. ForteBio Octet Data Analysis software was used to align and fit the data in 1:1 binding mode ($K_{on}$ = 7.29 x $10^4$ $M^{-1}s^{-1}$, $K_{off}$ = 8.27 x $10^{-3}$ $s^{-1}$ and $K_D$ = 1.14 x $10^{-7}$ M).

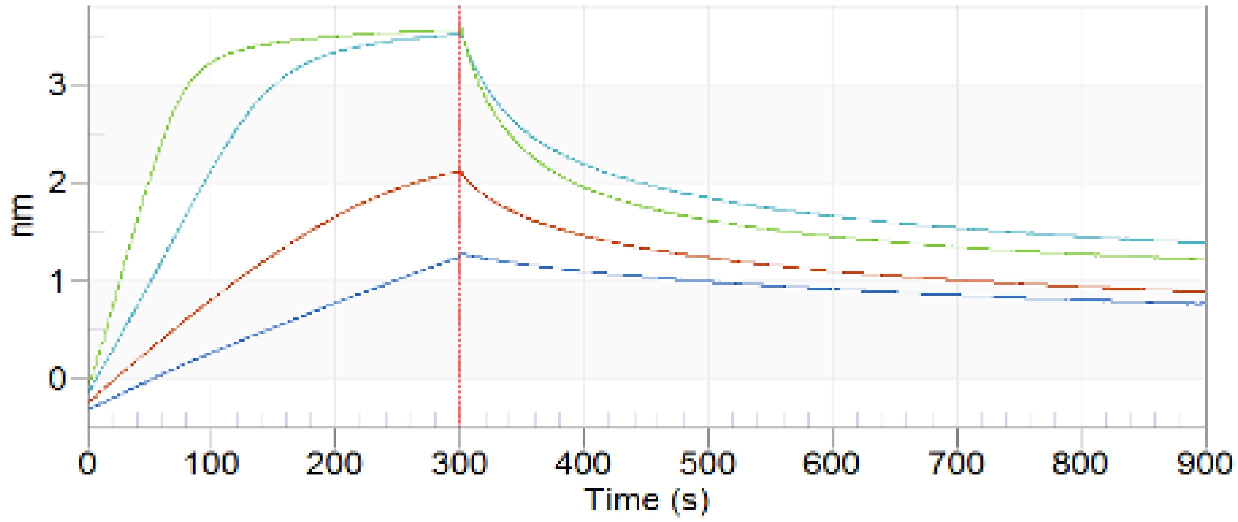


**Figure S6**. BLI affinity measurements of the TBA-based ISD. SA tips were pre-hydrated in ISD buffer for 20 minutes and then in 20 nM TBA to achieve 0.5 nm loading level. After ligand loading, sensors were dipped in 25, 50, 100, 200 nM thrombin for 300 sec and then dipped in buffer for 600 sec. Signals were normalized by subtraction of the signal from reference tips. ForteBio Octet Data Analysis software was used to align and fit the data in 1:1 binding mode ($K_{on}$ = 4.12 x $10^4$ $M^{-1}s^{-1}$, $K_{off}$ = 1.01 x $10^{-2}$ $s^{-1}$ and $K_D$ = 2.46 x $10^{-7}$ M).

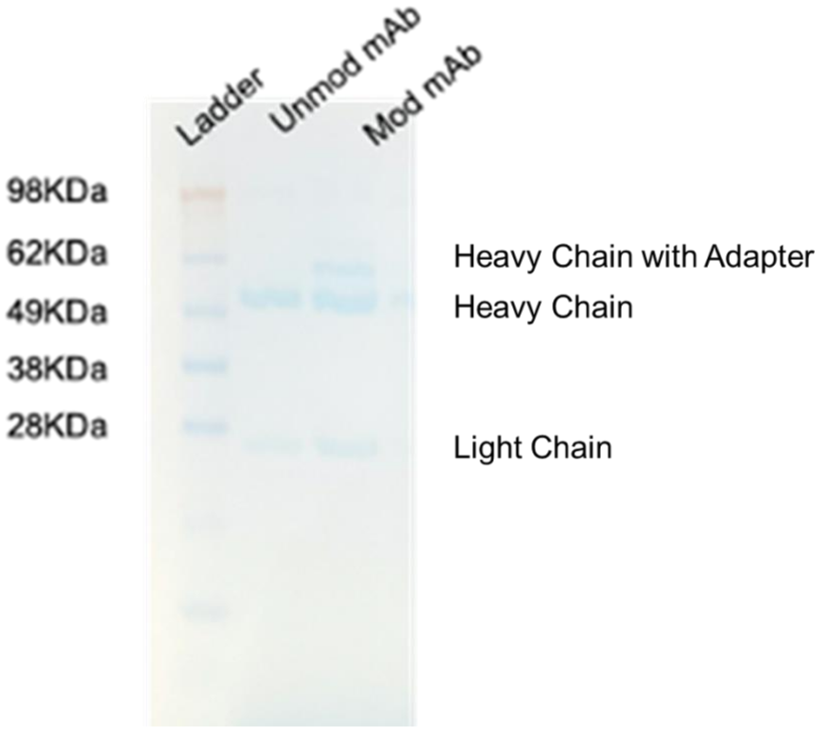


**Figure S7**. We assessed PANDAS antibody labeling via 10% Nu-PAGE. Lane 1 was loaded with SeeBlue ladder. Lane 2 was loaded with unmodified antibody 5020. Lane 3 was loaded with modified 5020 in preparation for PANDAS assembly. In Lane 3, the shifted heavy chain and unmodified heavy chain are present at a roughly 1:1 ratio, indicating that each mAb has one oligo tether on average.

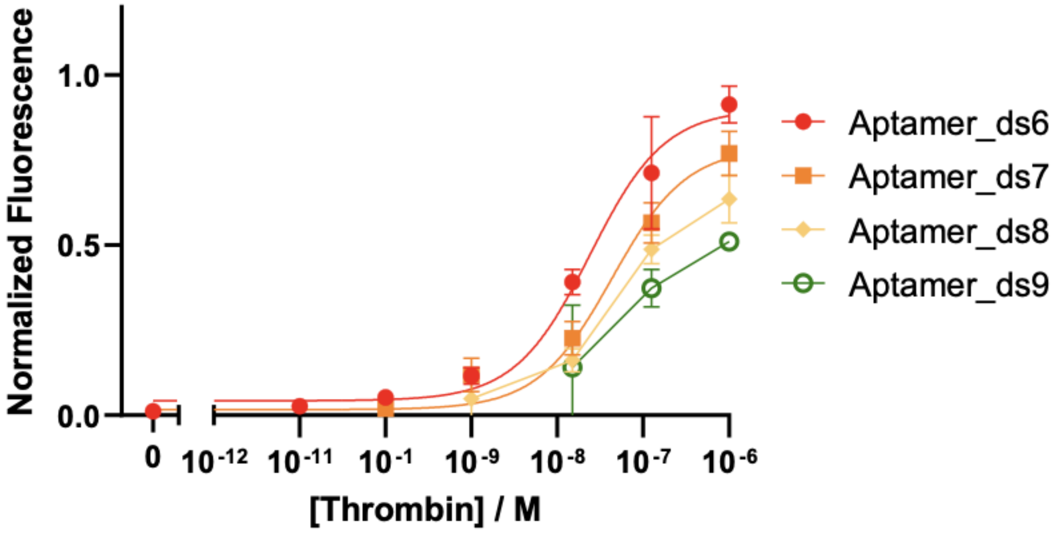


**Figure S8**. We assessed how tuning of DS length affected ISD performance. For consistency with our standard PANDAS assembly workflow and protein G bead-based read-out in flow cytometry, we assembled these ISDs with an antibody against TNF-α that showed no affinity for thrombin.

#### Supplementary Discussion: Thermodynamic binding model for the PANDAS construct

We begin with the assumption that the target concentration $T > U_i$, where $U_i$ is the total concentration of PANDAS in solution (initially in state $U$), and then assign the following Boltzmann weights in accordance with the equilibria described in **Figure 2a**:

| Partition | U | $B_{Ap}$ | $B_{Ab}$ | $B_{Ap,Ab}$ | $B_{Ap}B_{Ab}$ |
|---|---|---|---|---|---|
| Weight | 1 | $T/K_{D,Ap}$ | $2T/K_{D,Ab}$ | $2TC_{eff}/(K_{D,Ap}K_{D,Ab})$ | $2T^2/(K_{D,Ap}K_{D,Ab})$ |

We can then derive an expression for the fraction of fluorescent molecules as a function of target, $F(T)$, as:

$$F(T) = \frac{B_{Ap} + B_{ApAb} + B_{Ap}B_{Ab}}{U_i} = \frac{T\left(2C_{eff} + K_{DAb}\right) + 2T^2}{K_{DAp}K_{DAb} + T\left(2C_{eff} + K_{DAp} + 2K_{DAb}\right) + 2T^2}. \quad (Eq. S1)$$

Eq. S1 is plotted as the solid lines in **Figure 2**. If we assume that $K_{DAp}, C_{eff} \gg K_{DAb}$, as we can expect from a PANDAS employing a high affinity antibody and a short tether, Eq. S1 can be approximated as the sum of two scaled isotherms, and is easily related to the qualitative interpretation provided in the main text. The first isotherm has a high affinity, with an effective dissociation constant defined by:

$$K_{D,HA} = \frac{K_{DAp}K_{DAb}}{2(K_{DAp} + C_{eff})}, \qquad (Eq. S2)$$

which describes the antibody-assisted binding of the aptamer. The second isotherm has a low affinity, and is defined by $K_{DLA} = K_{DAp} + C_{eff}$, which describes the un-assisted binding fraction. Note that the scaling of the isotherms depends on the relative values of $K_{DAp}$ and $C_{eff}$. If $C_{eff} << K_{DAp}$, then most of the binding will be unassisted, while if $C_{eff} \gg K_{DAp}$, the majority of signal will be from antibody-assisted binding. Thus, we can define the parameter $\alpha = \frac{C_{eff}}{K_{DAp}+C_{eff}}$, which describes the fraction of antibody-assisted binding of a given PANDAS system, such that:

$$F(T) \approx \alpha \cdot \frac{T}{\alpha K_{DAp} K_{DAb} / 2C_{eff} + T} + (1 - \alpha) \cdot \frac{T}{(K_{DAp} + C_{eff}) + T}. \qquad (Eq. S3)$$

For the case where $C_{eff}$ is large, antibody-assisted binding dominates, and $\alpha \approx 1$. In this case, Eq. S3 approaches Eq. 1.
